# AptaBlocks: Accelerating the Design of RNA-based Drug Delivery Systems

**DOI:** 10.1101/216465

**Authors:** Yijie Wang, Jan Hoinka, Piotr Swiderski, Teresa M Przytycka

## Abstract

Synthetic RNA molecules are increasingly used to alter cellular functions. These successful applications indicate that RNA-based therapeutics might be able to target currently undruggable genes. However, to achieve this promise, an effective method for delivering therapeutic RNAs into specific cells is required. Recently, RNA aptamers emerged as promising delivery agents due to their ability of binding specific cell receptors. Crucially, these aptamers can frequently be internalized into the cells expressing these receptors on their surfaces. This property is leveraged in aptamer based drug delivery systems by combining such receptor-specific aptamers with a therapeutic “cargo” such that the aptamer facilitates the internalization of the cargo into the cell. The advancement of this technology however is contingent on an efficient method to produce stable molecular complexes that include specific aptamers and cargoes. A recently proposed experimental procedure for obtaining such complexes relies on conjugating the aptamer and the cargo with complementary RNA strands so that when such modified molecules are incubated together, the complementary RNA strands hybridize to form a double-stranded “sticky bridge” connecting the aptamer with its cargo. However, designing appropriate sticky bridge sequences guaranteeing the formation and stability of the complex while simultaneously not interfering with the aptamer or the cargo as well as not causing spurious aggregation of the molecules during incubation has proven highly challenging. To fill this gap, we developed AptaBlocks, a computational method to design sticky bridges to connect RNA-based molecules (blocks). AptaBlocks relies on a biophysically inspired theoretical model capturing the complex objectives of the design and yet is simple enough to allow for efficient parameter estimation. Given this model, the sticky bridge sequence is optimized using a Monte Carlo algorithm based on heat-bath transitions. The effectiveness of the algorithm has been verified computationally and experimentally. AptaBlocks can be used in variety of experimental settings and its preliminary version has already been leveraged to design an aptamer based delivery system for a cytotoxic drug targeting Pancreatic ductal adenocarcinoma cells. It is thus expected that AptaBlocks will play a substantial role in accelerating RNA-based drug delivery design.

AptaBlocks is available at https://github.com/wyjhxq/AptaBlocks.

## 1 Introduction

Synthetic RNA molecules are increasingly utilized to alter the behavior of genes and cells [1, 2, 3, 4]. For example, RNA interference (RNAi) can be used as a highly sequence-specific gene silencing mechanism. In addition, programmable RNAs can be engineered to enact and tune gene regulatory mechanisms through mediating interactions with the cellular machinery [2]. Furthermore, *cis*-acting RNAs are subject to regulation through interactions with *trans*-acting small RNAs (sRNAs) to enable or block their formation, creating inducible genetic control elements [2, 3]. RNA-based therapeutics hence hold a promise for targeting currently undruggable genes and treating a diverse array of diseases including cancer [5, 6]. Recently, RNA aptamers emerged as promising drug delivery agents (reviewed in [7, 8]). Aptamers are short RNA or DNA molecules identified through iterative rounds of *in vitro* selection [9, 10] to specifically recognize and bind cognate targets. Importantly, aptamers that bind to specific cell receptors can often be internalized into the cells that express these receptors on their surface [7] making them ideal candidates as cell specific vehicles to deliver therapeutic cargoes.

Initially, aptamer based delivery systems were assembled by conjugating aptamers with cargoes of interest through synthesis of long RNA molecules containing both sequences [11, 12, 13]. However, this strategy suffers from the considerable disadvantage of requiring the synthesis of long RNA molecules. Recently, Zhou, et al. [14, 15] proposed an alternative approach that relies on conjugating complementary RNA strands to the aptamer and the cargo to form aptamer-stick and cargo-stick conjugates respectively. When incubating the conjugates together in a subsequent step, these complementary strands would hybridize to form a double-stranded “sticky bridge” connecting the aptamer with its cargo (Fig. 1) [14, 15]. The experimental design hence follows a three-step procedure where aptamer-stick and cargo-stick are first synthesized and allowed to fold in separate buffers (Step 1-2, Fig. 1A-I and A-II), followed by an annealing phase in a binding buffer to form the final aptamer-sticky bridge-cargo complex (Fig. 1A-III). As described later in this paper, this procedure not only reduces the length of the sequences to be synthesized but, if designed appropriately, allows to reuse the aptamer-stick as a universal delivery agent for several different cargoes.

**Fig. 1:**
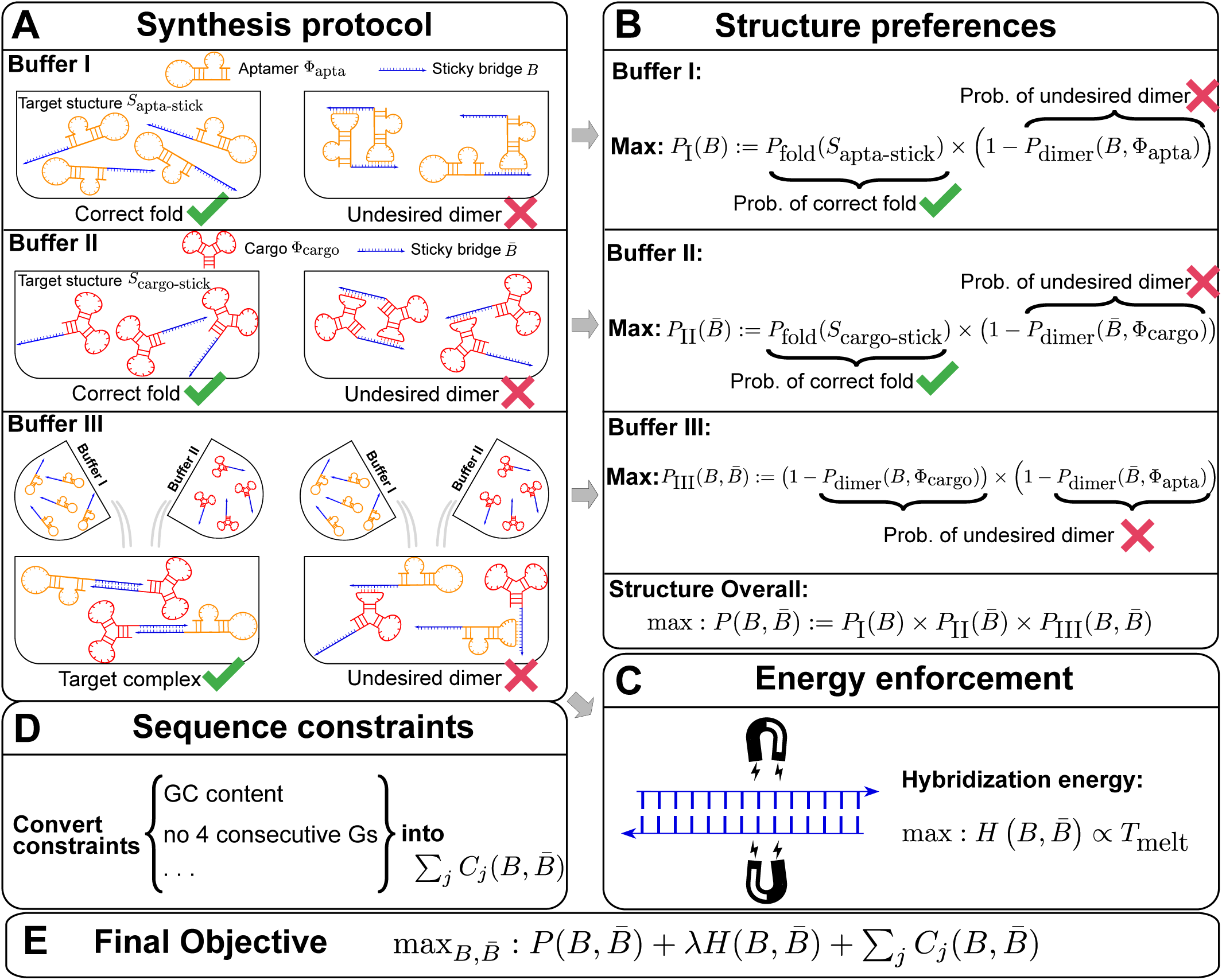
(A) The synthesis protocol [14] that AptaBlocks relies on. (A-I) Production of the first intermediate, corresponding to the conjugation *Φ*_apta-stick_(*B*) of the aptamer *Φ*_apta_ and the sticky bridge *B* folding into structure *S*_apta-stick_. Notably, *S*_apta-stick_ includes all structures where *B* is unpaired. (A-II) Production of the second intermediate, which is the the conjugation *Φ*_cargo-stick_(*S*) of the cargo *Φ*_cargo_ and the sticky bridge 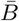 forming into structure *S*_cargo-stick_. Importantly, *S*_cargo-stick_ includes all structures where 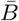 is unpaired. (A-III) Generation of the final RNA complex. (B) Converting structure preferences in each buffer into thermodynamic probabilities. (C) Enforcing strong binding between sticky bridges to increase melting temperature. 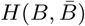) is defined in Eq. (5). (D) Formulation of sequence constraints into indicator functions. (E) Final objective function that incorporates structure preferences, energy enforcement and sequence constraints.

**Fig. 2.**
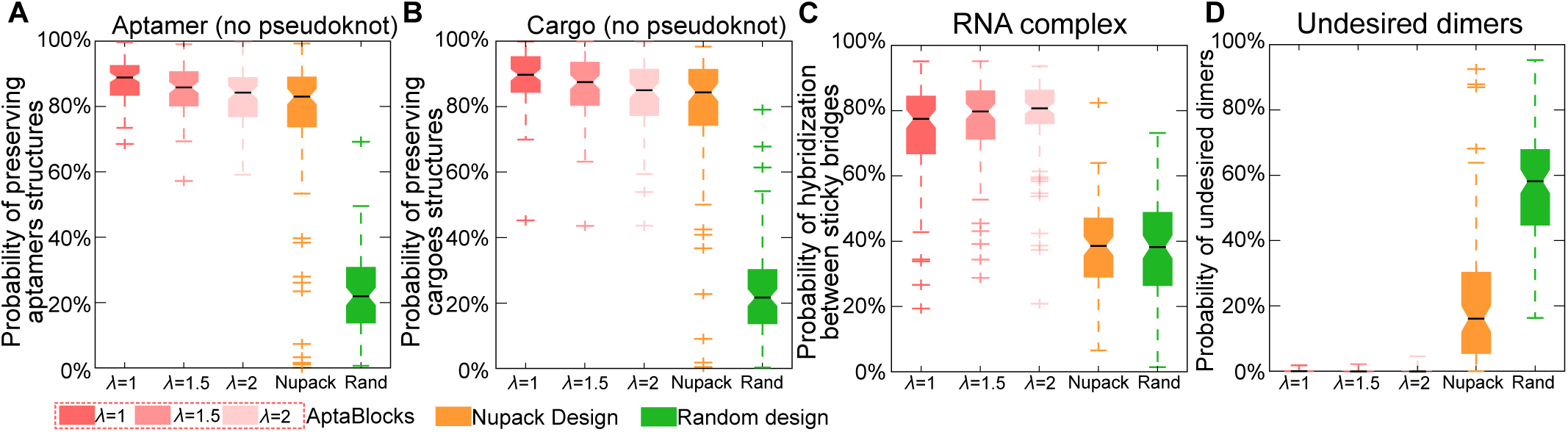
Comparison of the competing algorithms. (A) Comparison of conserving the secondary structure of aptamers. The probability of sticky bridges being unpaired is computed using a secondary structure model without pseudoknots. (B) Comparison of conserving the secondary structure of cargoes. The probability of sticky bridges being unpaired is computed using a secondary structure model without pseudoknots. (C) Comparison of binding affinity. The binding affinity is approximated by computing the probability of hybridization between sticky bridges using an RNA-RNA interaction model. (D) Comparison on incurring undesired dimers. The probability of undesired dimers is computed by our proposed model in Supplementary Materials B. Larger *λ* indicates that AptaBlocks focuses more on increasing the melting temperature of sticky bridges.

However, the current design process for sticky bridges tends to be solely informed by the experimentalists expertise and in practice, manual designs have shown only limited success. This observation can be contributed to three main challenges which emerge at different stages of the assembly. First, the covalently added “sticky bridge sequences” must hybridize to form a stable double helix. Second, they cannot interfere with any secondary and tertiary structural elements of the aptamer and the cargo and third, unwanted aggregation of the molecules, such as cargo sticky bridge interactions, must be avoided.

To address these challenges, we propose AptaBlocks, an efficient computational method for designing aptamer-sticky bridge-cargo complexes. Accounting for the three-step procedure [14, 15], we formulate the sticky bridge sequence design as an optimization problem utilizing an objective function which reflects the biophysical characteristics of the assembly process. specifically, we designed the objective function considering the equilibrium probabilities of the target structures over all possible structures of the aptamer-stick and cargo-stick, the probability of the interaction between the aptamer-stick and cargo-stick at equilibrium, the hybridization energy between the sticky bridge sequences, and additional sequence constraints including but not limited to the GC content. We further provide a simulated annealing algorithm that enables efficient estimation of the corresponding combinatorial optimization problem.

When tested on simulated data, AptaBlocks successfully preserved the structures of both, the aptamers and cargoes while achieving high binding affinity between the complementary strands of the sticky bridges. Next, we measured the performance of our approach to design universal sticky bridges for one aptamer and multiple distinct cargoes by varying the number and the sequence similarity of the latter. Our results indicate that AptaBlocks is capable of designing universal sticky bridges for either many cargoes with similar sequences or a smaller number of cargoes with distinct sequences. Finally, we experimentally validated our findings *in vitro* by successfully applying AptaBlocks to design sticky bridges for an aptamer-siRNA conjugate.

Despite our focus on ssRNA or dsRNAs as cargoes, it is important to note that our approach extends naturally to a multitude of other cargoes. As a case in point, a universal bridge designed with AptaBlocks is currently explored in an experimental setting, facilitating the conjugation of cytotoxic drugs and RNA aptamer tP19, targeting pancreatic tumor cell growth [16]. Given the rapid development of efficient *in vitro* selection technologies allowing for the generation of highly target-specific and target-affine aptamers, we anticipate that with its general design strategy, AptaBlocks will further accelerate the development of flexible and effective drug delivery systems.

## 2 Results

### 2.1 Method Overview

AptaBlocks takes two or more RNA molecules (the building blocks) as input and designs an RNA complex containing these molecules while preserving their individual secondary structures and thus their biological functions. In a typical application, one of the molecules is an RNA aptamer (a delivery agent), while the other molecule, the so called “cargo”, is a therapeutic with the potential of modifying the cell’s function after being internalized. Note that designing Aptamer-Cargo complexes is a special case of the problem of designing RNA complexes with particular properties [17, 18]. Indeed, our method is general and can be applied to more complex settings such as combining more than two RNA molecules. To facilitate such designs, AptaBlock does not require that the sticky bridge is necessarily at the termini of the assembled molecules. Furthermore, for designing an RNA complex that include a larger number RNA molecules more than one sticky bridge can be used.

In the case of two building blocks, an aptamer and a cargo, AptaBlocks aims at designing two complementary RNA sequences *B* and 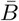, so that when *B* is conjugated with the aptamer and 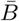 is conjugated with the cargo, the desired aptamer-cargo complex is successfully formed through hybridization between *B* and 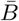.

Note that this model assumes a specific order of assembly as described in [14] where the building blocks are synthesized and folded separately and then incubated together to allow formation of the final complex (Fig. 1A). Consistent with this experimental set-up, AptaBlocks designs two intermediates, *Φ*_apta-stick_ and *Φ*_cargo-stick_, the former referring to the aptamer conjugated with *B* and the latter to the cargo conjugated with 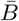 while satisfying the following conditions:

– *B* and 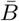 are unstructured in *Φ*_apta-stick_ and *Φ*_cargo-stick_ and, in particular, do not perturb the structures of the aptamer and the cargo. Examples of these target structures are shown in Fig. 1 A-I and A-II (target structure).
– Two or more *Φ*_apta-stick_ molecules should not bind to each other to form undesired dimers or larger aggregates. The same condition has to be satisfied by *Φ*_cargo-stick_. Examples of these undesired dimers are shown in Fig. 1 A-I and A-II.
– When *Φ*_apta-stick_ and *Φ*_cargo-stick_ incubate together, the sticky bridge strand *B* in *Φ*_apta-stick_ should not interact with the cargo strand in *Φ*_cargo-stick_. Similarly, the sticky bridge strand 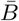 in *Φ*_cargo-stick_ should not interact with the aptamer strand in *Φ*_apta-stick_. Examples of these off-target interactions are shown in Fig. 1 A-III (undesired dimer).
– To maximize stability of the complex, the melting temperature of the sticky bridge that controls complex formation is maximized, i.e. the binding free energy between *B* and 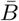 is minimized.

Additional constraints on the sticky bridge sequences, such as the exclusion of long chains of G’s due to their difficulty of being synthesized, can also be added. Given the complex interplay between the above mentioned objectives, a trade-off between optimizing one over another exists. Thus AptaBlocks generates several alternative solutions with their respective scores so that they can be further investigated if required.

In what follows, we describe our method in more details. We first outline our biophysics based model to compute the probabilities that characterize the structural properties mentioned above. Then, we elucidate the heat-bath algorithm, a Monte Carlo method closely related to the classical (Metropolis-based) simulated annealing, that AptaBlocks uses to optimize these probabilities as well as the melting temperature, subject to possible additional constraints the user might impose on *B* and 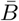.

### 2.2 The AptaBlocks Algorithm

In this section we formulate the optimization function allowing us to design sticky bridges with the properties outlined in the previous subsection.

**Preserving the original functionality of interacting RNA molecules** The function of a molecule is dependent on both - its sequence and structure. Therefore conjugating *B* with the aptamer and 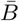 with the cargo respectively can affect the desired and functionally active structures of the aptamer and cargo. Thus the aptamer-stick conjugate *Φ*_apta-stick_(*B*) is required to attain a specific target structure *S*_apta-stick_ that consists of the aptamer part folded in the same manner as before it was conjugated with *B* while ensuring that *B* does not interact with the aptamer and remains unfolded (Fig. 1 A-I). Preserving the target structure therefore translates into maximizing the probability *P*_fold_ (*S*_apta-stick_|*Φ*_apta-stick_ (*B*)) [19, 20] that the sequence *Φ*_apta-stick_ (*B*) folds into *S*_apta-stick_. To achieve this, it suffices to enforce all nucleotides of *B* to be unpaired in *Φ*_apta-stick_(*B*). Recall that in the experimental protocol, *Φ*_apta-stick_ (*B*) molecules are initially present in one buffer, say Buffer I, opening the possibility of binding to each other (Fig. 1 A-I). Assuming that the aptamers themselves do not dimerize, dimerization between the *Φ*_apta-stick_(*B*) molecules at the sticky bridge strand *B* must still be prevented. Denoting *P*_dimer_ (*B*, *Φ*_apta_) as the probability of undesired binding between the *B* strand of one molecule with the *Φ*_apta_ part in another, we can formulate the structural preferences mentioned above for *Φ*_apta-stick_ (*B*) as Eq. (1) (Fig. 1B for Buffer I) and further maximize Eq. (1) to secure structural properties for *Φ*_apta-stick_ (*B*).

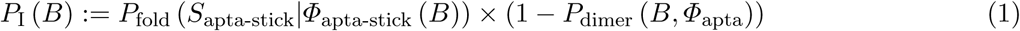

Similarly, to ensure the cargo-stick species in Buffer II fold into the target structure *S*_cargo-stick_ which preserves the fold of the cargo and restricts 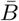 to be unpaired, as well as to prevent the formation of undesired dimerizations, we maximize 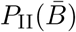 defined as (Fig. 1B for Buffer II)

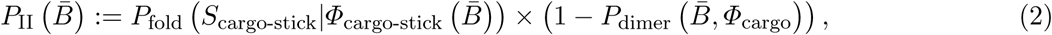

where 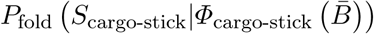 is the probability that the cargo-stick species folds into *S*_cargo-stick_ for given 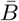 [19, 20], and 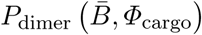 corresponds to the dimerization probability between 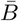 and *Φ*_cargo_ across species. In case of a double stranded cargo such as a siRNA, 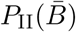 is computed differently from a single-stranded RNA molecule and we refer the reader to Supplementary Material A for a detailed discussion.

To generate the final RNA complex, the above two intermediates are mixed together in a third buffer, expecting the complementary sticky bridge strands *B* and 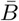 to hybridize together. To avoid undesired interactions between sticky bridge strands and loop-structures (functional components [21, 22]) of the aptamer and the cargo, we reduce the probability of those off-target dimerizations (as shown in Fig. 1 A-III) by maximizing 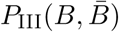 (Fig. 1B for Buffer III):

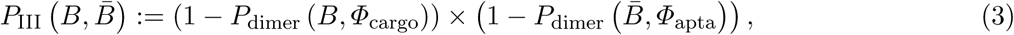

where *P*_dimer_ (*B*, *Φ*_cargo_) is the dimerization probability between sticky bridge *B* and cargo strand *Φ*_cargo_ and 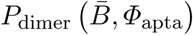 stands for the dimerization probability between sticky bridge 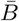 and aptamer strand *Φ*_apta_.

Overall, the sticky bridge strands *B* and 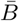 have to simultaneously maximize 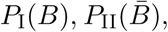 and 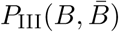 to guarantee the production of two intermediates and the final aptamer-sticky bridge-cargo RNA complex. Hence, we define the overall structure objective 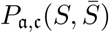 for aptamer a and cargo c as

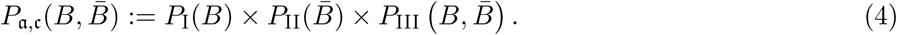

**Optimizing the melting temperature** To increase the melting temperature between *B* and 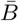, which is proportional to the free energy of their hybridization [23], we minimize the hybridization free energy between the sticky bridge sequences *B* and 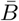 by maximizing the energy objective function defined as

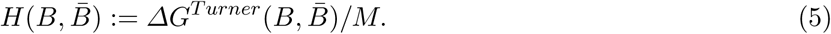

where the hybridization energy 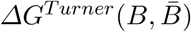 is computed based on the Turner RNA folding model [24]. *M* is used to normalize 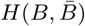 into [0, 1], which can be set as the smallest 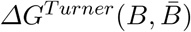.

**Sequence constraints for sticky bridges** Besides conforming to structural preferences and optimizing melting temperature, further specific sequence requirements might have to be satisfied. In addition to ensuring complementarity between *B* and 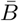, one might for instance require a particular GC content or must take into account chemical synthesis constraints prohibiting the inclusion of 4 consecutive guanines in the primary structure. Assuming *C* is a sequence constraint, we define an indicator function *C* as

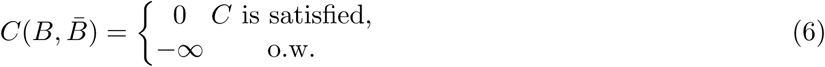

for inclusion into the final model as outlined below.

**Final formulation** Considering the structure preferences, energy enforcement and sequence constraints, we formulate the sticky bridge design problem as

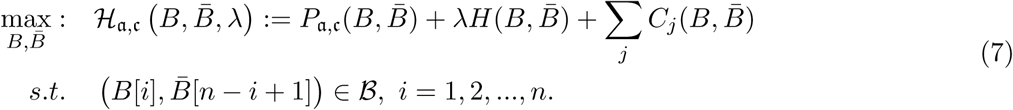

where the constraint enforces *B* and 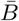 to be complementary and 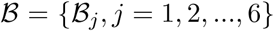 corresponds to a set of all possible 6 base pairs with 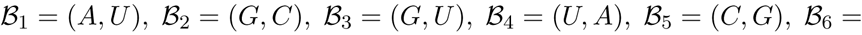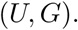 The length of the sticky bridge sequences is defined as 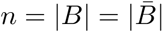 whereas *C*_*j*_ corresponds to the *j*th sequence constraint.

To take advantage of the complementary property between *B* and 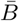, we define 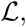, where 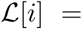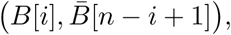 and therefore transform the problem of designing sequences *B* and 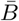 into designing base pairs for 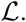 After abuse of the notations, we convert problem (7) into

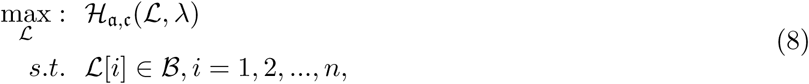

**Extension to multiple cargoes setting** In practical applications, the goal is often to target a specific cell using an appropriate aptamer and deliver a cargo optimized to manipulate specific pathways. If, for instance, we aim at silencing a gene using siRNA, we are typically not limited to a single siRNA species to perform this task. In addition, if we are not successful in silencing a gene we might try to silence another gene in the same pathway. However, synthesizing a different *B* and 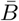 for each possible aptamer-cargo pair is highly impractical, time consuming, and cost intensive. Notably, our flexible model enables to bypass this issue by extending our approach to allow for designing a *universal sticky bridge* for a specific aptamer but multiple different cargoes. Let a be an aptamer of interest and ℭ be a set of cargoes that need to be delivered. Then we can attempt to find the optimal universal stick bridge sequences for all aptamer-cargo combinations by solving

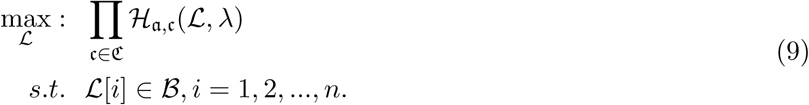

**The optimization procedure** To optimize our target function, subject to all imposed constraints, we use a heat-bath Monte Carlo optimization strategy - a strategy similar to the classical simulated annealing. While both, simulated annealing and the heat-bath, are conceptually very similar, the classical simulated annealing is based on the Metropolis approach to compute transition probabilities without computing partition functions which is computationally intractable in many applications. In our setting however, efficient computation of the partition function is not a challenge, allowing us to use the alternative heat-bath transitions probabilities which are known to have better performance [25, 26].

Let 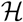 represent the Hamiltonian, the objective function of the combinatorial problem. We apply the single spin heat-bath update rule, which updates the system energy when making a base pair change for a given position *i* in 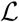 from base pair 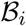 to base pair 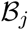 while keeping the rest of the base pairs in 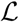 fixed: 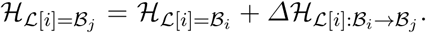 The probability of making the base pair change is proportional to the exponential of the corresponding energy change of the entire system, i.e

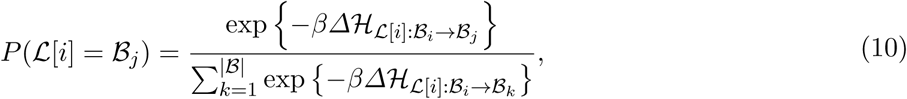

where *β* = 1*/T*. The details of the AptaBlocks algorithm are elaborated in Algorithm 1.

#### Algorithm 1

The AptaBlocks algorithm. Here, *T*_*start*_ and *T*_*end*_ are the start and end temperatures, *c* stands for the cool down procedure, and *K* is the number of sweep time for each temperature. *I* = {1, 2, …, *n*} is a set containing the position indices of the sticky bridge 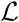 and *λ* is the parameter in (8) and (9) that balances between meeting the structure preferences and increasing the melting temperature.

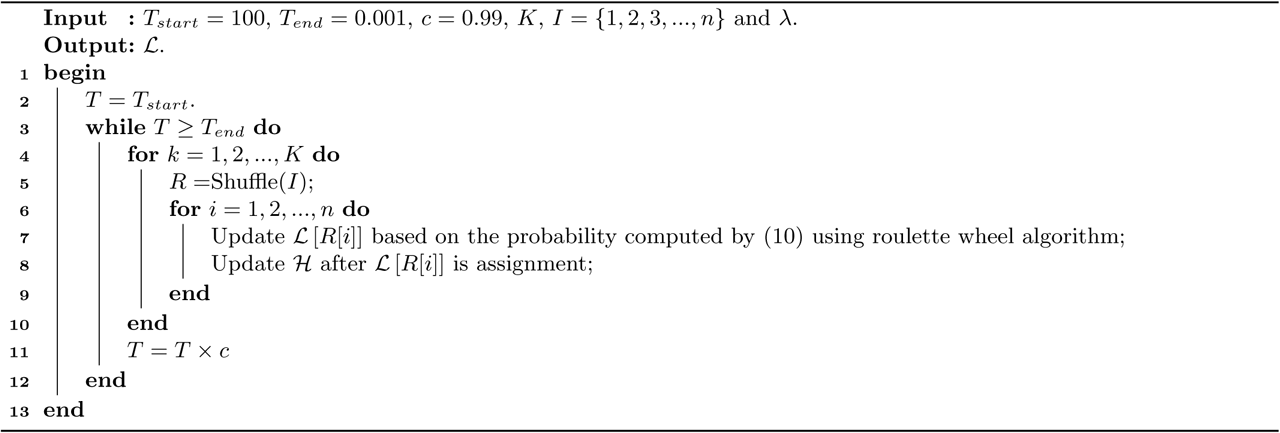

**Efficient approximation** Solving problems (8) and (9) using the AptaBlocks algorithm is computationally expensive. Based on Eq. (10), in the AptaBlocks algorithm **Algorithm 1**, the calculation of the Hamiltonian 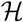 is a fundamental operation and is repeatedly executed. The time complexity of computing 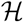 is dominated by calculating the dimerization probability *P*_dimer_(·, ·) required in Eqs. (1), (2), and (3). *P*_dimer_(·, ·) can be computed based on an RNA-RNA interaction model [27, 28] therefore defining the time complexity of computing 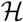 as *O*(*m*^6^), where *m* = max(|*Φ*_apta_|, |*Φ*_cargo_|).

To reduce the time complexity, we propose to prevent undesired dimerization through minimizing 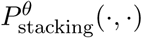 instead of *P*_dimer_(·, ·). 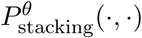 is the probability of strong stacking hybridization between pairwise interacting RNAs where the binding free energy is less than *θ*. Based on Lemma 1 in Supplementary Materials B, we prove

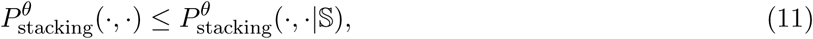

where 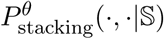 is the probability of strong stacking hybridization with free energy less than *θ* conditioned on that the pairwise RNAs indeed bind together only through stacking hybridization. Therefore, computing and minimizing 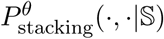 the upper bound of 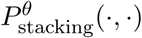 can eventually minimize 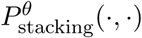 to avoid strong binding across species.

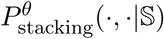 can be computed in *O*(*m*^3^) based on the model developed in Supplementary Materials B. In summary, we replace *P*_dimer_(·, ·) in Eqs. (1), (2), and (3) by 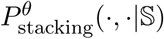 allowing the time complexity of computing 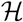 to be reduced to *O*(*m*^3^).

### 2.3 Results on Simulation Data

**Designing sticky bridges for single aptamer-cargo pairs** As the first step towards validation of our approach we applied AptaBlocks to simulated data. We generated 100 aptamer sequences and 100 ssRNA cargo sequences at random. Next, we randomly combined these sequences and obtained 100 aptamer-cargo pairs. Finally, we set the length of the sticky bridge for each aptamer-cargo pair. To balance the difficulties of these 100 design tasks, we used the same aptamer length for all 100 tasks. Similarly, we set the same length for all cargoes and sticky bridges, respectively. The parameters used to generating the simulation data are available at Supplementary Materials C.1.

We applied AptaBlocks, Nupack Design [17], and an approach in which sticky bridges were obtained as random complementary sequences of the given length to the simulation dataset and compared the quality of the results obtained with each method. Notably, to the best of our knowledge, Nupack Design is the only existing algorithm that in theory could design sequences for sticky bridges. specifically, Nupack Design strives to obtain RNA complexes by concatenating interacting RNA strands as a single strand and enforcing the sticky bridge to be formed on the single strand.

Performance was determined by testing whether the original structures of aptamers and cargoes are preserved, whether the binding affinity of the sticky bridges is strong, and whether undesired dimerizations are present. Hence, we measured the preservation of the original structures of aptamers by computing the probabilities that the sticky bridges stay unpaired using both a secondary structure model without pseudo-knots [19, 20, 29] and a secondary structure model with pseudoknots [30, 31]. We stress that our structure optimization procedure did not include pseudoknots due to the prohibitive cost of such computations. Similarly, we determined the preservation of the structures of cargoes by computing the probabilities that the sticky bridges stay unpaired in cargo-stick conjugates using both the secondary structure models with and without pseudoknots. The binding affinity of sticky bridges can be estimated by the probability that the sticky bridge strands hybridize across aptamer-stick and cargo-stick conjugates using an RNA-RNA interaction model [27, 28]. Probability of undesired dimerizations is estimated by our stacking hybridization model introduced in Supplementary Materials B. We designed 10 sticky bridges for each aptamer-cargo pair and used the average of the above probabilities in comparison.

The comparisons between AptaBlocks and other approaches are detailed in Fig. 1. Figs. 1A and B show that AptaBlocks outperforms other approaches on preserving structures of aptamers and cargoes. Despite that AptaBlocks’ design does not take pseudoknots into account, it performed quite well on the energy model with pseudoknots Fig. S2, again significantly outperforming its competitors. Furthermore, we observe that the sticky bridges designed by AptaBlocks have higher probabilities to hybridize than other designs by a large margin as shown in Fig. 1C, but lower probabilities to incur undesired dimerizations as shown in Fig. 1D.

Next, we tested the impact of the length of the sticky bridge on each method. In this setting, we use the same simulation data but assign different lengths of the sticky bridges and analyze the performance of preserving the structures of aptamers, cargoes, and binding affinity of the sticky bridges as the function of the length of the sticky bridges.

We applied AptaBlocks and Nupack Design to the simulation data and compared their performance in terms of the probabilities of preserving original structures of aptamers and cargoes, the probability of hybridization between sticky bridges, and the probability of incurring undesired dimerizations. For each aptamer-cargo pair, we run each competing method 10 times and report the resulting averages. The overall averages of the 100 aptamer-cargo pairs are shown in Fig. 3 whereas the experimental details can be found in Supplementary Materials C.2.

**Fig. 3:**
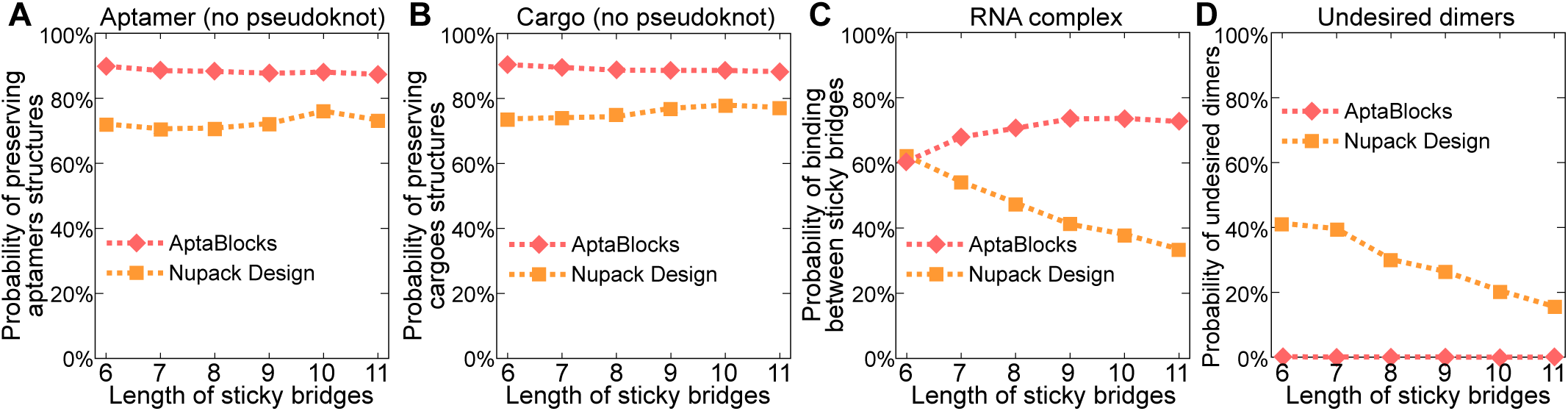
Comparison of the competing algorithms on different length of sticky bridges. (A) Comparison of conserving the secondary structure of aptamers for different length of sticky bridges. The probability of sticky bridges being unpaired is computed using a secondary structure model without pseudoknots. (B) Comparison of conserving the secondary structure of cargoes for different length of sticky bridges. The probability of sticky bridges being unpaired is computed using a secondary structure model without pseudoknots. (C) Comparison of binding affinity for different length of sticky bridges. The binding affinity is approximated by computing the probability of binding between sticky bridges using an RNA-RNA interaction model. (D) Comparison on incurring undesired dimers. The probability of undesired dimers is computed by our proposed model in Supplementary Materials B.

Fig. 3 illustrates the comparison between AptaBlocks and Nupack Design. Fig. 3 A and B indicate that AptaBlocks are more capable of preserving the structures of aptamers and cargoes for the sticky bridges with increasing length than Nupack Design. Fig. 3 C shows that AptaBlocks achieves better binding affinity than Nupack Design, which loses control of binding between sticky bridges when the length of the sticky bridges increases. In contrast, we find that the longer the sticky bridges are the better the binding affinity AptaBlocks can achieve. After a certain length however, the binding affinity remains constant, suggesting that we can determine the optimal length of the sticky bridges by observing the changes of the binding affinity for AptaBlocks as shown in Fig. 3 C. Fig. 3 D illustrates that comparing to Nupack Design the sticky bridges designed by AptaBlocks rarely lead to undesired dimerizations.

**Designing universal sticky bridges** Using the same 100 simulated aptamer-cargo pairs as in the previous section, we further analyzed AptaBlocks performance on the task of designing universal sticky bridges for a single aptamer and multiple alternative cargoes. For this, we selected the cargo sequence as the seed for each aptamer-cargo pair and generated additional cargo sequences by mutating the seed at random. We used sequence similarity, the percentage of identical bases between sequences, to measure similarity between cargo sequences and made sure that the pairwise similarities between all pairs of cargo sequences in the same group are identical. The generation of the simulation data is elaborated in Supplementary Materials C.3.

Analogous to the previous sections, we then evaluated the performance by computing the probabilities of preserving the original structures of aptamers and cargoes, as well as the probabilities of hybridization between sticky bridges. For each design, we run AptaBlocks and Nupack Design 10 times and computed the average over those probabilities. For more details regarding the implementation of this experiment, we refer the reader to Supplementary Materials C.3. Notably, Nupack Design [17] was not originally designed for this task but can be used to generate sticky sequences by treating the aptamer-stick and each cargo-sticks as reactants and each aptamer-sticky bridge-cargo species as an intermediate [17].

Fig. 4 depicts the comparison between AptaBlocks and Nupack Design on designing universal sticky bridges. Fig. 4 A and B show that AptaBlocks outperforms Nupack Design by a large margin. In addition, Fig. 4 A implies that the original structures of cargoes and aptamers become hard to preserve when the number of cargoes get larger and the cargo sequences are more distinct. Similarly, as shown in Fig. 4B, the binding affinity between universal sticky bridges weakens with increasing number of cargoes and distance across cargo sequences. Overall, the experiment suggests that AptaBlocks can design promising universal sticky bridges for a large number of cargoes of similar sequences or a small number of cargoes with very different sequences.

**Fig. 4:**
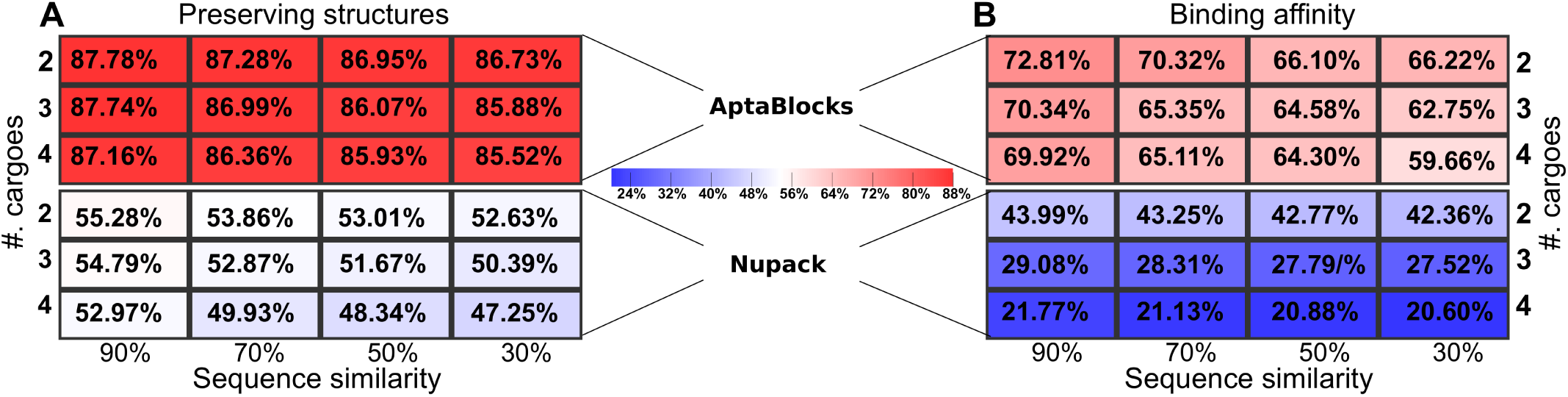
Quality of universal sticky bridge design of AptaBlocks and Nupack Design for various number of cargoes and sequence similarity between cargoes. (A) Probabilities of preserving the original structures of the aptamers and cargoes (based on the secondary structure model without pseudoknots) for different number of cargoes and different sequence similarity between cargoes. (B) Probabilities of binding between the designed universal sticky bridges for different number of cargoes and different sequence similarity between cargoes. Note that Nupack Design was not originally designed for this purpose and while it can be used to perform this task as described in the text, it does not ensure high binding affinity nor structural preservation.

### 2.4 Experimental Validation

Complementing the *in silico* validation of AptaBlocks, we also validated its ability to design sticky bridges *in vitro* on a pancreatic cancer cell model using aptamer tP19 as described in [16] and a siRNA as shown in Fig. 5A. We explored the design of combining the aptamer and the single siRNA cargo. The resulting molecules as suggested by AptaBlocks were consequently synthesized and allowed to form the final complex in accordance with steps ①-④ in Fig. 5A. The successful formation of each molecule and/or conjugate was monitored by running an agarose gel for each step (Fig. 5B). The bands verify the presence of the complex without noticeable undesired dimerizations.

**Fig. 5:**
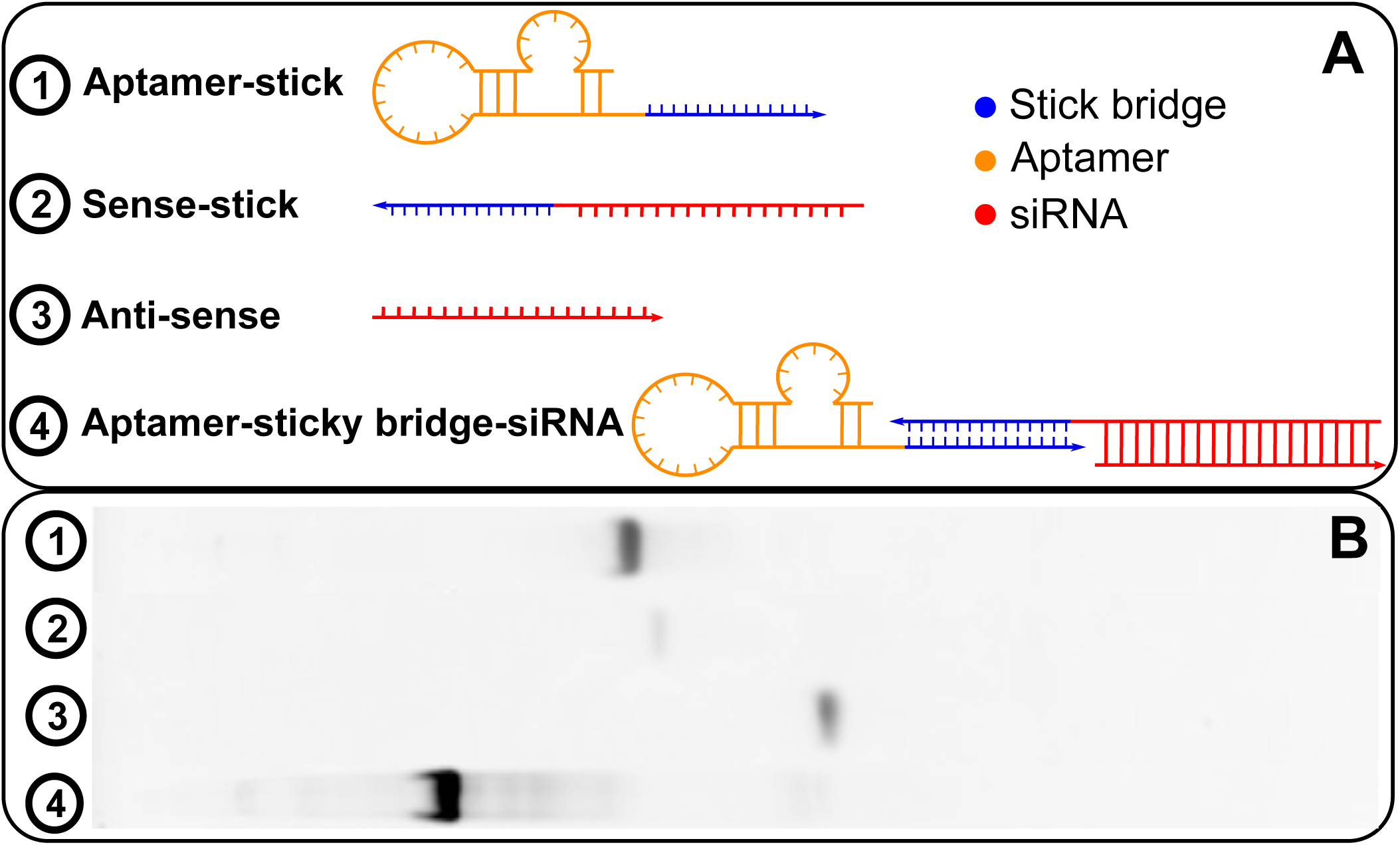
(A) The RNA molecules of aptamer-stick, sense-stick, anti-sense, and the RNA complex of aptamer-sticky bridge-siRNA. (B) The agarose gel image for all species showing the successful formation of all species without undesired dimerizations.

In the same pancreatic cancer model, but in contrast to conjugating a siRNA to the aptamer, AptaBlocks has also been successful in designing a sticky bridge for an aptamer-drug conjugate. This design has been implemented experimentally, verified in vitro, and described in detail in [16].

## 3 Discussion

Aptamer based drug delivery systems provide unparalleled opportunities for targeted drug therapies mainly due to the ability of designing aptamers to recognize specific cells and thus delivering intervention only to these cells. Indeed, the aptamer-drug conjugates designed using our AptaBlocks strategy reported in [16] were shown to significantly inhibit cell proliferation of pancreatic cancer cells in a dose-dependent manner while at the same time showing minimal cytotoxicity in normal cells and in the control MCF7 cell lines.

Synthesizing long conjugates containing a concatenation of an aptamer and a cargo has several drawbacks and thus a more flexible method based on non-covalent conjugation using a sticky bridge has been recently proposed [14, 15]. While improving the experimental procedure, the method hinges on the successful design of the sticky bridges connecting aptamers and cargoes. This design task initially relied on an expensive and time consuming try and error approach and its success was highly dependent on the experimentalists expertise. This challenge, together with a high failure rate, showed that creating sticky bridges for a diverse array of applications could greatly benefit from a computationally informed design approach.

Until now, to the best of our knowledge, only Nupack Design [30, 17] could be adopted to tackle the problem. However, Nupack Design [17] was not able to consistently provide a satisfying solution even for the simplest variant of the problem that considers only one aptamer and one cargo. The main reason was that Nupack Design is making assumptions that are not consistent with our experimental setting. Nupack Design concatenated input molecules which, in the context of our application, has lead to spurious designs. In addition Nupack Design ignores the interactions between different RNA strands that might cause undesired dimerization which is the most frequent reason for designs failing *in vitro*.

Thanks to the carefully designed theoretical model, AptaBlocks has proven to be successful in computational and experimental settings. It not only incorporates structural constraints required for a correct synthesis, but also optimizes hybridization strength of complementary strands in the sticky bridge, and minimizes formations of undesired dimerization and higher order aggregations. While in this paper, we focus on two key applications of AptaBlocks, designing a sticky bridge for two RNA molecules (an aptamer and a cargo) and designing a universal sticky bridge allowing for conjugating the same aptamer with several alternative cargoes, the method is general and can applied to other RNA complex designs such as the ones shown in Fig. 5(A).Given the broad utility of the approach in the context of RNA-based drug delivery systems and its success in a variety of experimental settings, AptaBlocks has the potential to greatly accelerate research efforts in this area.

## 4 Acknowledgement

The authors would like to thank Sorah Yoon for providing the agarose gel for experimental validation.

## Supplementary Materials

### A Structure preferences for double stranded RNA cargoes

AptaBlocks can also design sticky bridges for dsRNA cargoes, such as siRNA. The structure preferences in Buffer II and Buffer III are different to the ones introduced in the main manuscript. Let *Φ*_sense_ and *Φ*_anti_ represent the sense and anti-sense parts of an siRNA. Assume the sticky bridge 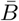 is conjugated to the sense part *Φ*_sense_ of the siRNA and let 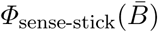 be the conjugation between *Φ*_sense_ and 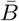 In Buffer II, we expect structure *S*_sense-stick_ (shown in Fig. S1 Buffer II) be the dominant structure for 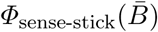 and therefore the probability 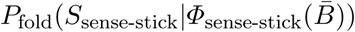 should be maximized. At the same time, undesired dimerizations between the sticky bridge 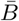 and either the sense part *Φ*_sense_ of the siRNA or anti-sense part *Φ*_anti_ of the siRNA should be avoid. Considering all those preferences, we maximize

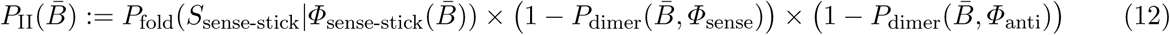

**Fig. S1.**
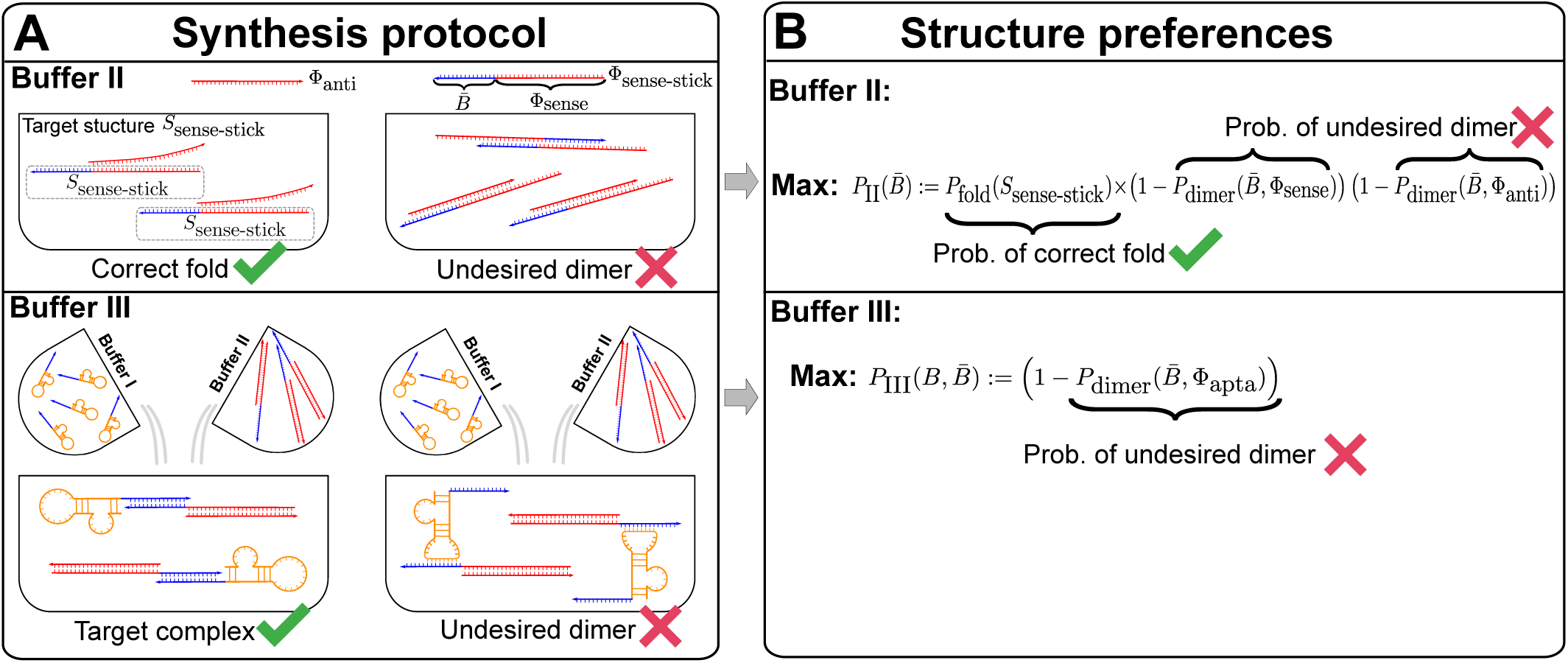
Structure preferences in Buffer II and III for dsRNA cargoes. (A) The target structure and undesired dimer in Buffer II and III. (B) Converting structure preferences in (A) into to objective functions.

For Buffer III, we only need to prevent dimerzation between the sticky bridge 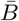 and the aptamer *Φ*_apta_.

Hence, we define 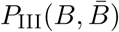 as

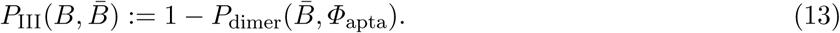

### B Stacking hybridization model for RNA-RNA interaction

In this paper, to reduce time complexity but still achieve reasonable results, we propose to minimize 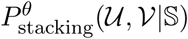 instead of 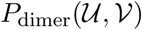 (time complexity *O*(|V|^6^)) [25, 26]) to prevent dimerization between 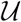 and 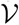. In fact, we are not interested in computing the exact thermodynamic probability 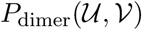 that 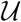 and 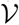 dimerize. Instead, our goal is to avoid strong dimerization between 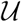 and 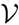. Hence, we propose to reduce the probability 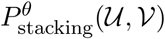 that 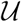 and 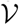 interact by stacking hybridization with free energy less than *θ*. Based on the following lemma, we know that minimizing 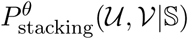 the upper bound of 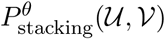 can eventually force 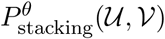 to decrease. Here, 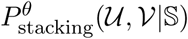 is the probability that 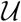 and 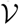 interact by stacking hybridization with free energy less than *θ* conditioned on that 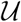 and 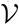 are actually interact only by stacking hybridization.

##### Lemma 1. 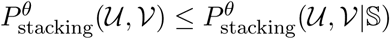

*Proof*. We define ℂ to represent all conformations that 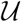 and 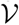 interact by stacking hybridization with free energy less than *θ* and 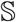 to represent all conformations that 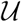 and 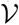 interact by stacking hybridization. Based on the definitions, we know 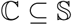. Then we have

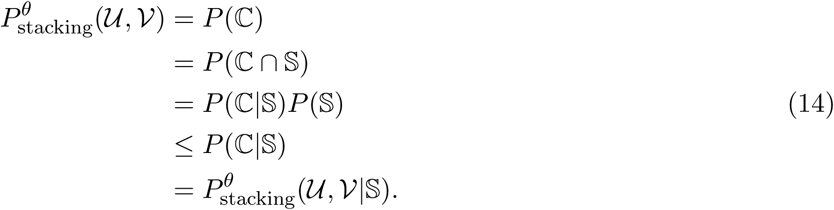

The third line in Eq. (14) is derived based on Kolmogorovs definition. The fourth line in Eq. (14) is derived based on 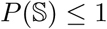.

The probability 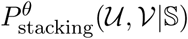 can be efficiently computed as following. Let *U* and *V* be the sequences of two RNA molecules and assume |*U*| < |*V*|. The free energy of the stacking hybridization of length *w* between 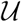 staring at *U*_*k*_ and 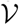 staring from *V*_*k'*_ is *ΔG*^*s*^(*U*_*k*_, *V*_*k'*_, *w*). The energetics of RNA-RNA interactions can be viewed as a stepwise process, therefore, *ΔG*^*s*^(*U*_*k*_, *V*_*k'*_, *w*) consists of the energy for exposing the binding site in the appropriate conformation for individual molecules and the energy gain from hybridization at the binding site [21]. Then the partition function of stacking hybridization between 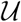 and 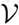 can be calculated by

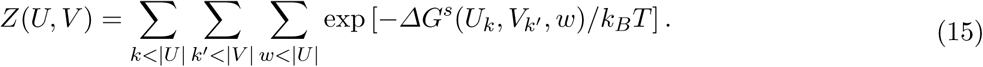

Based on the same framework, the partition function of stacking hybridization with free energy smaller than *θ* can be computed by

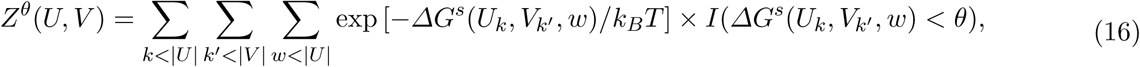

where *I*(*a* < *b*), an indicator function, equals 1 when *a* < *b* and 0 otherwise. Both *Z*^*θ*^(*U*, *V*) and *Z*(*U*, *V*) can be computed in *O*(|*V*|^3^). Then we can compute the probability 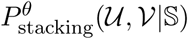 that 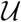 by (17) in *O*(|*U*|^2^|*V*|).

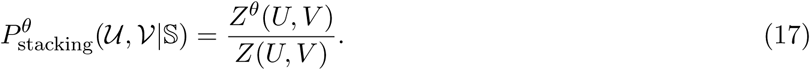

### C Implementation details

#### C.1 Designing sticky bridges for single aptamer-cargo pair

We first generated 100 aptamer sequences of size 10 drawn from a uniform nucleotide distribution. In the same fashion, we generated 100 ssRNA cargo sequences of size 10. Next, we randomly combined those 100 aptamer and 100 cargo to obtain 100 aptamer-cargo pairs. For each aptamer-cargo pair, we set the length of the sticky bridge to 10 base pairs. Two sequence strands in the sticky bridges are respectively concatenated to aptamers and cargoes. The design scenario is similar to Fig. 1.

Other than the length of the sticky bridge, AptaBlocks and Nupack Design need one more parameter, respectively. AptaBlocks needs *λ* to balance between structure preferences and energy enforcement. We assigned *λ* = 1.0, 1.5, 2.0. Nupack Design needs a random seed for initialization.

For each simulated aptamer-cargo pair, we ran AptaBlocks 10 times for each *λ* and ran Nupack Design 10 times using a different random seed. Then we computed the averages of the probabilities of preserving the structures of the aptamers and the cargoes, the probabilities of hybridization between the designed sticky bridges, and the probabilities of incurring undesired dimers. The average of these results is summarized and detailed in Fig. 2.

#### C.2 Length of sticky bridges

We utilized the same simulation dataset generated in C.1 but we varied the length of the sticky bridges from 6 base pairs to 11 base pairs. For each aptamer-cargo pair, we ran AptaBlocks 10 times using *λ* = 1.0 and Nupack Design 10 times using 10 different random seeds. We computed the averages of the probabilities of preserving the structures of the aptamers and the cargoes, the probabilities of hybridization between the designed sticky bridges, and the probabilities of incurring undesired dimers over the 10 designs. We associated the average probabilities to each aptamer-cargo pair and display the averages over 100 aptamer-cargo pairs in Fig. 3.

#### C.3 Designing universal sticky bridges

We designed universal sticky bridges for one aptamer and multiple ssRNA cargoes using AptaBlocks. We utilized the same simulation dataset generated in C.1, but for each aptamer-cargo pair, we used the cargo sequence as the seed to generate additional cargoes with of identical size. For this, we first, set all cargo sequences to the seed cargo sequence. Then, we randomly selected several locations on the sequences according to the desired sequence similarity and mutated that position. Here, sequence similarity is defined as the ratio of the number of matching bases between two sequences to the length of the sequences. Furthermore, for cargoes that need to be delivered by the same aptamer, we ensure that the pairwise sequence similarity between any two cargoes is the same.

For each design, we ran AptaBlock 10 times using *λ* = 1.0 and Nupack Design 10 times with 10 random seeds, and computed the average performance. For a total of 100 designs, we computed the average over the average performance for each design and report the result in Fig. 5. Notably, we computed the probabilities of preserving the original structures rather than computing the probabilities for aptamers and cargoes separately as Fig.4.

### D Comparison based on the secondary structure model with pseudoknots

We compared AptaBlocks with Nupack Design on preserving structures of aptamers and cargoes using a secondary structure model with pseudoknots. The following figures shows their performance. Both AptaBlocks and Nupack Design do not consider pseudoknots in their designs. But Nupack Design used the same energy parameters used by Nupack Analysis which we used to compute the probability of preserving structures considering pseudoknots. Although Nupack Design has an advantage by using the same energy parameters, AptaBlocks still outperforms this method.

**Fig. S2.**
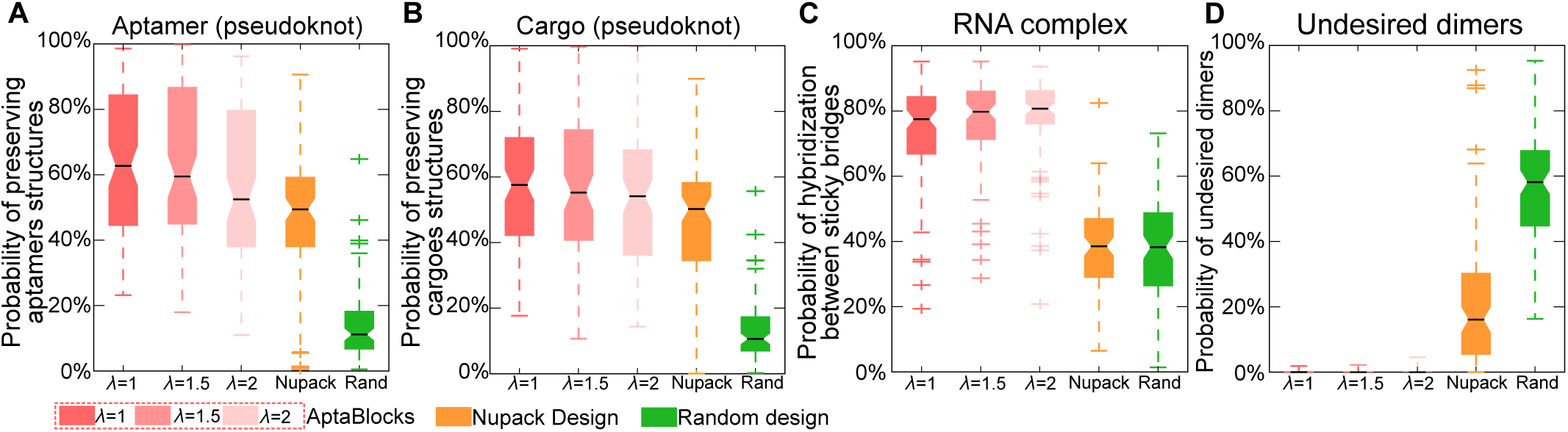
Comparison of the competing algorithms. (A) Comparison on conserving the secondary structure of aptamers. The probability of sticky bridges being unpaired is computed using a secondary structure model with pseudoknots. (B) Comparison on conserving the secondary structure of cargoes. The probability of sticky bridges being unpaired is computed using a secondary structure model with pseudoknots. (C) Comparison on binding affinity. The binding affinity is approximated by computing the probability of hybridization between sticky bridges using RNA-RNA interaction model without pseudoknots. Comparison on incurring undesired dimers. The probability of undesired dimers is computed by our proposed model in supplementary materials B. Larger *λ* indicates that AptaBlocks focuses more on minimizing binding free energy between sticky bridges.

